# Measuring mtDNA turnover, synthesis, and supercoiling via selective bromodeoxyuridine incorporation

**DOI:** 10.1101/2023.10.24.563889

**Authors:** Jingti Deng, Armaan Mohan, Timothy E Shutt

## Abstract

An often-overlooked aspect of mitochondrial biology is the mitochondrial DNA (mtDNA). The multi-copy mtDNA is highly dynamic, with changes in supercoiling, synthesis rates, and turnover rates that are tightly associated with mitochondrial and cellular functions. To better understand the state of mtDNA, here we describe a protocol that selectively incorporates bromodeoxyuridine into mtDNA for subsequent measurement via an adapted Southern blot followed by immunoblotting (a.k.a. Southwestern blot). This basic protocol can be applied with slight modifications for the measurement of mtDNA synthesis, turnover or supercoiling to understand mtDNA changes.

## 1. Introduction

Mitochondria harbour their own genome (mitochondrial DNA or mtDNA), a remnant of their bacterial ancestry. In humans, the circular 16 569 bp mtDNA genome encodes 37 canonical genes, which include 13 proteins, two ribosomal RNAs and 22 tRNAs [1,2]. Unlike the nuclear genome, mtDNA is present in hundreds of copies per cell and replicates independently of the cell cycle in a manner that is tightly regulated. The importance of properly maintaining mtDNA is evidenced by changes to mtDNA copy number associated with disease [3]. Given the circular nature of mtDNA, supercoiling is another important aspect that can affect the genome function [4]. Understanding how mtDNA is regulated is crucial for understanding mitochondrial function [5,6]. As a result, there is a need for methods that monitor changes to mtDNA topology and replication. Incorporation of the thymine analog 5’-bromo-2’-deoxyuridine (BrdU) into nuclear DNA is a frequently applied technique to measure cell proliferation and cell cycle progression [7–10]. BrdU incorporation during DNA synthesis can be measured by antibodies against BrdU via immunohistochemistry, flow cytometry or other antibody-based methods [11,12]. While BrdU incorporation is typically used for monitoring nuclear DNA synthesis, this approach can also be used to monitor mitochondrial DNA (mtDNA) synthesis and turnover [13–15,6,16–23], as well as changes in mtDNA topology (*i.e.*, supercoiling) [24–26]. However, there are no detailed protocols available for using BrdU incorporation to measure these dynamic aspects of mtDNA. Here we describe how to use BrdU to label mtDNA, which can then be visualized following agarose gel electrophoresis and an adapted Southern transfer and immunoblot protocol. As mtDNA is far less abundant than nuclear DNA, approaches to minimize nuclear DNA labeling, such as the nuclear DNA synthesis inhibitor aphidicolin [27], or enrich mtDNA may be required. Overall, this BrdU incorporation approach, using common instruments and reagents, can be applied with slight modifications to measure multiple aspects of mtDNA dynamics including rates of mtDNA replication (Figure 1), mtDNA turnover (Figure 2), and mtDNA supercoiling (Figure 3).

**Figure 1:**
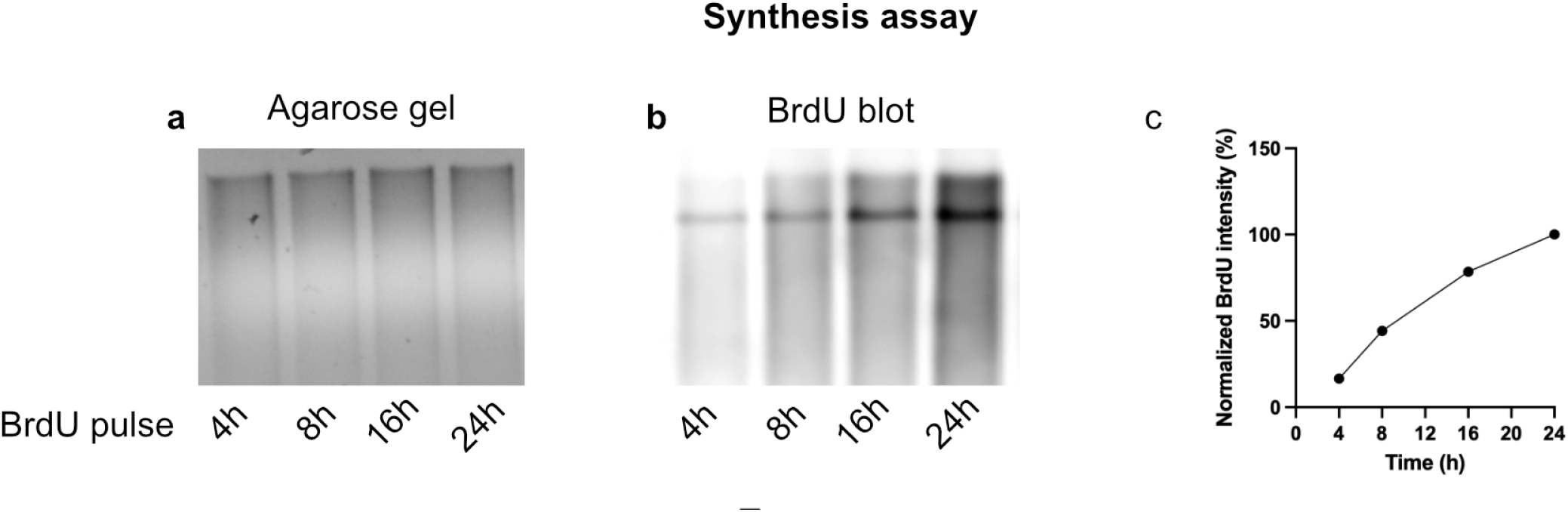
Representative results from mtDNA synthesis assay. 0.5 µg of total DNA isolated from U2OS cells treated as indicated was separated on agarose gels (**a**) and BrdU visualized after transfer and immunoblotting (**b**). For the mtDNA synthesis assay, BrdU was incorporated for 4 h, 8 h, 16 h and 24 h prior to DNA isolation. Total DNA was digested using *Bam*HI. The BrdU signal from the synthesis assay was quantified via ImageJ densiometry (**c**). Raw intensity values were plotted as percentage values normalized to 24 h post-BrdU incorporation.

**Figure 2:**
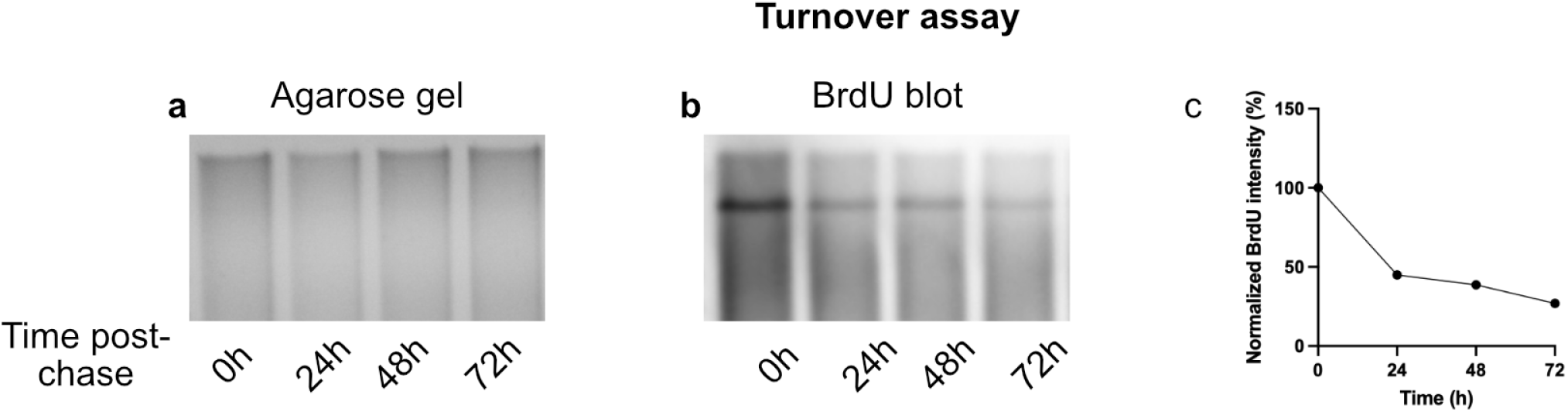
Representative results of a mtDNA turnover assay. 0.5 µg of total DNA isolated from U2OS cells treated as indicated was separated on agarose gels (**a**) and BrdU visualized after transfer and immunoblotting (**b**). For the mtDNA synthesis assay, BrdU was incorporated for 24 h prior to DNA isolation immediately (0 h) or 24 h, 48 h, 72 h post-chase. Total DNA was digested using *Bam*HI. The BrdU signal from the synthesis assay was quantified via ImageJ densiometry (**c**). Raw intensity values were plotted as percentage values normalized to 0 h post-chase.

**Figure 3:**
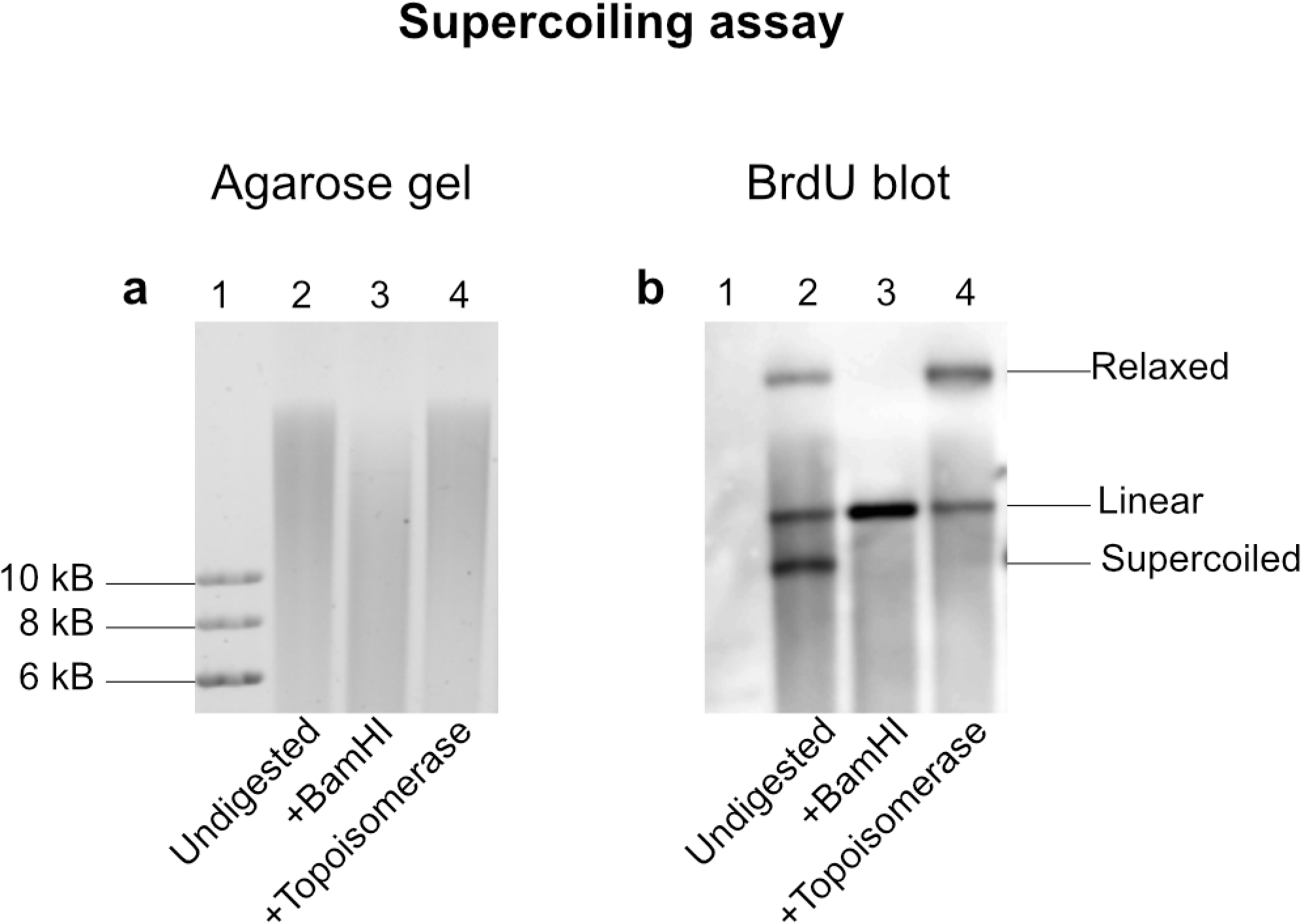
Representative results from the mtDNA supercoiling assay. 1 µg of total DNA isolated from HCT116 cells treated as indicated was separated on an agarose gel (**a**) and BrdU visualized after transfer and immunoblotting (**b**). The use of controls reveals the location of possible mtDNA topologies, as *Bam*HI digestion shows the location of the linear band, while treatment with topoisomerase removed the supercoiled band illustrating its location.

## 2. Materials

Prepare all solutions using ultrapure water that is DNAse-free, or sterile DMSO as noted. Ensure all solutions used on cultured cells are sterilized using a 0.22 µM filter, and all buffers are autoclaved. Storage conditions are listed for each solution.

### 2.1 Cell treatment

1. Cultured cells and cell culture medium suitable for the used cell type.
2. 6-well cell culture plates (35 mm diameter).
3. 1X PBS: 137 mM NaCl, 2,7 mM KCl, 10 mM Na_2_HPO_4_, 1.8 mM KH_2_PO_4_, pH 7.4. Sterilize by autoclaving and store at room temperature.
4. 0.25 % Trypsin/EDTA. Store at −20 °C and warm to 37 °C before use.
5. Aphidicolin (1000X): 10 mM in DMSO. Aliquot and store at −20°C (*see* **Note 1**).
6. 5’-Bromo-2’-Deoxyuridine (BrdU) (1000X): 50 mM in DMSO (1000X). Aliquot and store at −20 °C (*see* **Note 2**).
7. Uridine (1000X): 50 mg/mL in water. Store and aliquot at −20 °C.
8. Total DNA isolation kit (e..g., E.Z.N.A Tissue DNA kit, Omega Biotek).
9. Restriction enzyme with appropriate buffer to linearize mtDNA (e.g., for human mtDNA *Bam*HI).
10. Spectrophotometer (e.g., Nanodrop) or other method to quantify DNA concentration and purity.
11. Optional for topology analysis: Topoisomerase I enzyme

### 2.2 Agarose gel electrophoresis and Southern transfer

1. Horizontal electrophoresis system (gel size ca. 9 × 11 cm).
2. Agarose.
3. Tris-acetate-EDTA electrophoresis buffer (TAE; 50X): 2 M Tris, 1 M acetic acid, 10 mM EDTA in water. Store at room temperature.
4. Optional for topology analysis: Tris-borate-EDTA electrophoresis buffer (TBE; 10X): 1 M Tris, 0.9 M boric acid, 10 mM EDTA in water. Store at room temperature.
5. 10X DNA loading dye: 30 % glycerol, 1 mM EDTA, 0,25 % bromophenol blue, 0.25 % xylene cyanol. Store at room temperature.
6. DNA dye suitable for agarose gel electrophoresis or post-electrophoresis staining (e.g. Ethidium bromide, SybrSafe or GelRed)
7. Imaging system for visualization of the DNA in-gel (e.g., Chemidoc or similar)
8. Glass dish fitting the gel with its tray.
9. Saline-sodium citrate (SSC; 20X): 3 M sodium chloride, 0.3 M sodium citrate in water. Store at room temperature.
10. Denaturing buffer: 0.5 M sodium hydroxide, 1.5 M sodium chloride. Store at room temperature.
11. PVDF membranes (0.2 µM pore size)
12. Shallow container larger than the agarose gel (*e.g.*, a 9 in × 9 in Pyrex dish for a 9 cm × 11 cm gel)
13. Plastic platform accommodating the gel (e.g., the gel tray of the electrophoresis system)

### 2.3 BrdU Immunodetection

1. Tris-buffered saline stock (10X TBS): 250 mM Tris, 27 mM KCl, 1.37M NaCl, pH 7.4.
2. TBST: 1X TBS containing 0.1 % Tween-20.
3. TBSTM: 1X TBS containing 0.1 % Tween-20 and 5 % milk.
4. Mouse anti-BrdU antibody.
5. Anti-mouse HRP-linked secondary antibody.
6. High-sensitivity chemiluminescent HRP substrate (e.g., SuperSignal^TM^ West Femto Maximum Sensitivity Substrate, Thermo Scientific^TM^).
7. Filter paper (e.g., Whatman^TM^ grade 3MM cellulose paper).
8. Stack of paper towels.
9. Glass container (*e.g.*, square 9 in × 9 in Pyrex glass dish).
10. Weight of ca. 500 g (e.g., two 96-tube microcentrifuge racks weighing approximately 250g each).
11. UV crosslinker (e.g., a UV Stratalinker^TM^ 1800 or similar device, alternatively UV table).
12. Dish accommodating the membrane.

## 3. Methods

### 3.1 Cell culture and drug treatment

1. Seed cells at a high density in a 6-well plate (*see* **Note 3**) and allow cells to grow to 95 % confluency, approximately 2-3 days (*see* **Note 4**).
2. To block nuclear DNA replication prior to BrdU treatment, remove media and replace with fresh media containing 10 µM aphidicolin for at least four hours (*see* **Note 5**).
3. To initiate specific incorporation of BrdU into mtDNA, add BrdU (1000X stock) to a final concentration of 50 µM. For incubation times, refer to the specific protocols 3.6-3.8.

### 3.2 DNA isolation

1. Harvest cells via trypsinization. Remove media and add a sufficient volume of PBS to cover the well (approximately 1-2 mL). Aspirate PBS before adding a sufficient volume of 0.25 % trypsin-EDTA to cover the cells (approximately 300 µL for a 35mm well) and incubate at 37 °C until cells detach from the plate, about 1-2 minutes.
2. Add approximately 1 mL of complete cell culture media and transfer media containing trypsinized cells to a centrifuge tube. Pellet cells by centrifuging at 225 g for 5 minutes.
3. Rinse the cell pellet by re-suspending the cells in 2-5 mL of PBS before centrifuging at 225 g for 5 minutes to re-pellet the cells.
4. Remove PBS and immediately isolate total DNA from the cell pellet using a commercial kit and the manufacturer’s directions. Elute DNA in the minimum elution volume (*see* **Note 6**), and store at −20°C if necessary.
5. Measure DNA concentration, A260/280, and A260/230 using a Nanodrop spectrophotometer or similar. A260/280 and A260/230 should be approximately 1.8 and 2.0 respectively. A typical concentration is approximately 250-500 ng/µL.

### 3.3 Agarose gel electrophoresis

1. Optional restriction digest (see protocol 3.8): Digest 0.5 µg of total DNA with a restriction enzyme to linearize mtDNA and reduce nuclear background. For human mtDNA, *Bam*HI is recommended (*see* **Note 7**). Follow the manufacturer’s directions for composition and conditions of the restriction reaction.
2. Prepare a ca. 9 cm × 11 cm 0.45 % agarose gel in TAE buffer with 1.5 mm thick wells. This will require about 100 mL of agarose solution and yield wells with approximately 50 µL of loading volume and a 1 cm thick gel.
3. Place the gel into the electrophoresis chamber and cover with 1x TAE buffer
4. Add 10X DNA loading dye to the DNA samples to a final concentration of 1X and load DNA samples (*see* **Note 8**).
5. Run agarose gel at 55 V for 4 h at room temperature.
6. Stain the gel with a DNA dye (e.g., Ethidium bromide or SybrSafe (*see* **Note 9**).
7. Image agarose gel using an imaging system.

### 3.4 Transfer to PVDF membrane

1. Place the gel with its tray into a suitable dish. To minimize buffers for subsequent steps, the gel can be trimmed to approximately the top ¼ based on the migration of the ladder.
2. Rinse the gel briefly with water (approximately 40 mL) to remove the buffer.
3. Soak the gel in a sufficient volume of denaturing buffer to submerge the gel (approximately 40 mL) for 1 hour with gentle shaking.
4. Rinse the gel briefly with water to remove buffer.
5. Soak the gel in 10X SSC (approximately 40 mL) for 30 minutes.
6. While the gel is soaking, prepare the transfer platform. Fill a shallow container approximately halfway with 10X SSC (requires approximately 500 mL of buffer). Place a plastic platform large enough to support the agarose gel which elevates it sufficiently above the 10X SSC buffer (*e.g.*, the inverted casting tray for the agarose gel) (*see* **Note 10**). Cut and place a piece filter paper on top of the plastic platform large enough to overhang, touching the 10X SSC buffer (Figure 4).
7. Once the gel has been soaked, place it on top of the filter paper on the transfer platform which has been pre-wetted with 10X SSC.
8. Rinse a PVDF membrane of the size of the gel in 50 mL of ethanol for 10 s to activate it.
9. Layer the PVDF membrane on the gel. Pre-wet filter paper with 10X SSC and layer it on the membrane. Place a stack of dry paper towels at least 8 cm tall on top. Since the mtDNA bands are in the top half of the agarose gel when run using the above suggested runtimes, the gel and/or the PVDF membrane can be cut to only transfer in the top half.
10. Weigh down the paper towels with a light weight (*e.g.* two 96 microcentrifuge tube racks, weighing a total of approximately 500 g) on top of a plastic sheet protector and allow the gel to transfer via capillary action overnight at least 16 h. For the weight, any object heavy enough to compress the paper towels such that they are all in direct contact with each other will work.

**Figure 4:**
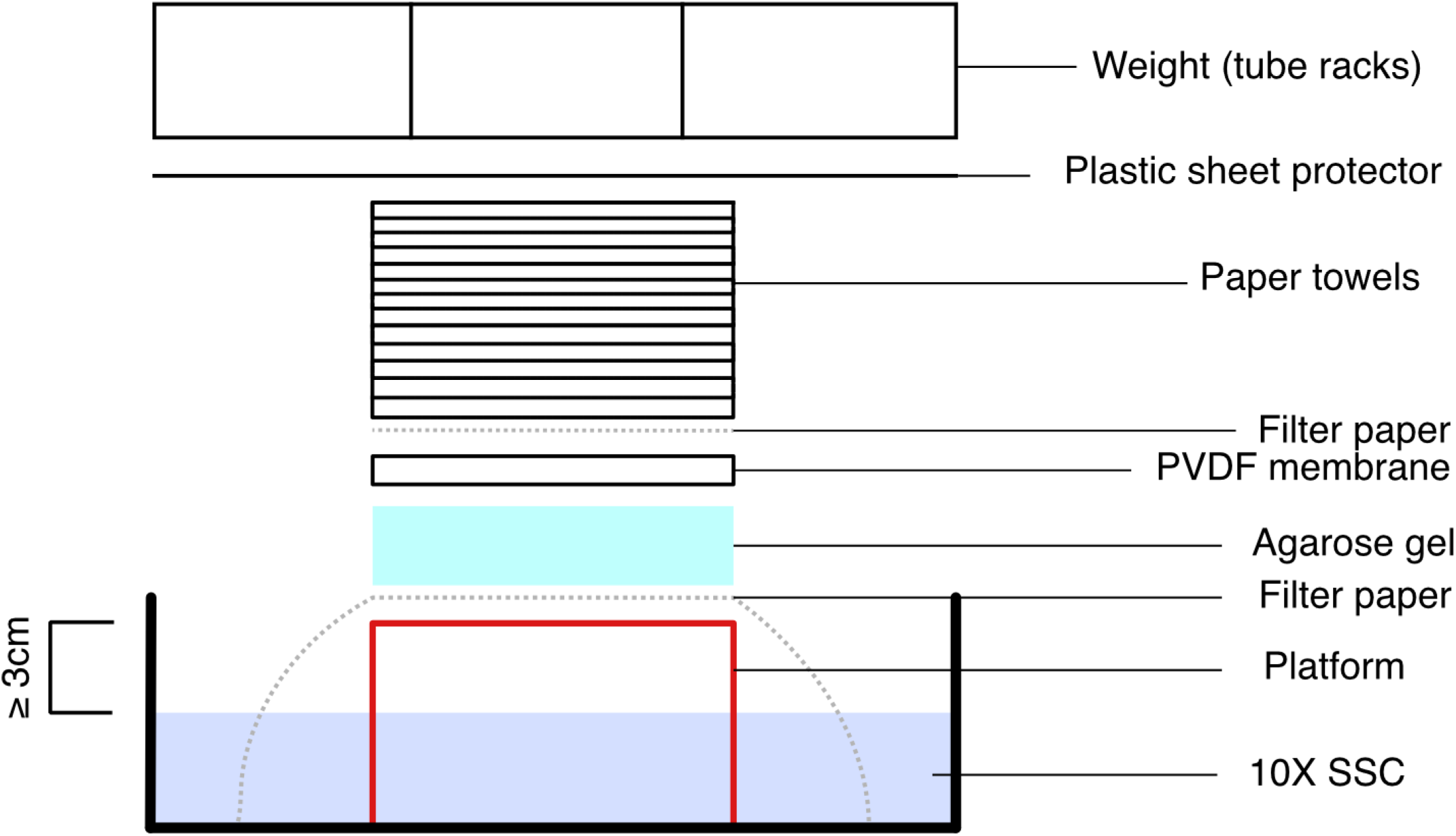
Diagram for assembly of the platform to transfer DNA from the agarose gel to the PVDF membrane. The agarose gel and PVDF membrane are sandwiched between a filter paper in contact with buffer and a dry filter paper, paper towels, and a weight system to allow the buffer to transfer DNA fragments by capillary action. The transfer is done at room temperature overnight.

### 3.5 BrdU immunodetection

1. Disassemble the transfer platform and soak the PVDF membrane in approximately 40 mL of 6X SSC for 5 minutes.
2. Place the PVDF membrane on a wet filter paper from the transfer stack to ensure that the membrane does not dry out.
3. Place it into a cross-linker and UV cross-link the PVDF membrane. Use the “auto-cross link” setting on a UV Stratalinker^TM^ 1800 or equivalent function on other machines.
4. Place the membrane into a dish and block it in TBSTM (approximately 15 mL) for 1 hour at room temperature.
5. Incubate membrane for 2 hours with anti-BrdU antibody (1:2000 in TBSTM, approximately 15 mL) at room temperature.
6. Wash membrane 3x for 5 minutes with TBST (approximately 15 mL each wash).
7. Incubate membrane with HRP-conjugated anti-mouse IgG (1:5000 in TBSTM, approximately 15 mL) for 1 hour.
8. Wash membrane 3x for 5 minutes for TBST (approximately 15 mL each wash).
9. Develop blots using ECL solution and image using a chemiluminescence imager (*see* **Note 11**). It is recommended to use a highly sensitive substrate such as SuperSignal^TM^ West Femto enhanced chemiluminescence substrate.

### 3.6 mtDNA synthesis assay (Figure 1)

1. Follow steps 1 and 2 of section 3.1, up to the the BrdU incorporation.
2. To observe the rate of mtDNA synthesis, perform BrdU incorporation over a range of timepoints. After treating with aphidicolin for at least 4 hours, add BrdU (1000X stock) to a final concentration of 50 µM. Incubate cells for desired timepoints (*e.g.*, 4, 8, 16, 24 hours).
3. Proceed immediately to isolate DNA as described under 3.2 (*see* **Note 12**).
4. Linearize mtDNA by performing a restriction enzyme digest as described under 3.3
5. Proceed with steps 3.3–3.5 and image the final BrdU immunoblot (Figure 1b).

### 3.7 mtDNA turnover assay (Figure 2)

1. Follow steps 1 and 2 in section 3.1, up to the BrdU incorporation.
2. For pulse-labelling of mtDNA while minimizing nuclear DNA incorporation, replace aphidicolin-only media with media containing 50 µM BrdU and 10 µM aphidicolin. Incubate cells for 24 h.
3. To measure mtDNA turnover, aspirate media and replace with fresh media containing 10 µM aphidicolin and 2 mM uridine (*see* **Note 13**). Incubate cells for the desired timepoints (*e.g.*, 24, 48, 72, 96 h) to allow previously BrdU-labeled mtDNA to be turned over.
4. Isolate DNA as described under 3.2. For step 3 of the DNA isolation, we recommend approaches to reduce nuclear DNA (*see* **Note 5**). Given the ratio of nuclear DNA to mtDNA, even low levels of BrdU incorporation into the nuclear genome can mask the mtDNA signal, as undigested nuclear DNA runs as a band of ~20 kb that can overlap with the region of interest for the mtDNA genome (Figure 5). In our hands, the following adaptations to the E.Z.N.A Tissue DNA kit greatly reduce the abundance of nuclear DNA. 1) Start with >2×10^7^ cells resuspend in 200 µL PBS. It is essential to use more than the recommended amount of cells. After addition of the BL lysis buffer, the solution should be viscous (this is critical for removal of nuclear DNA). 2) Before continuing with the DNA isolation procedure, run the viscous solution through a homogenizer column (*e.g.*, Omega Biotek, HCR 003) to reduce the viscosity (in this step, larger nuclear DNA is fragmented resulting in a homogeneous smear that does not interfere with the mtDNA signal). 3) Collect the flow through containing mtDNA and proceed with DNA isolation.
5. Linearize mtDNA by performing a restriction enzyme digest as described under 3.3.
6. Proceed with sections 3.3–3.5 and image the final BrdU immunoblot (Figure 2b).

**Figure 5:**
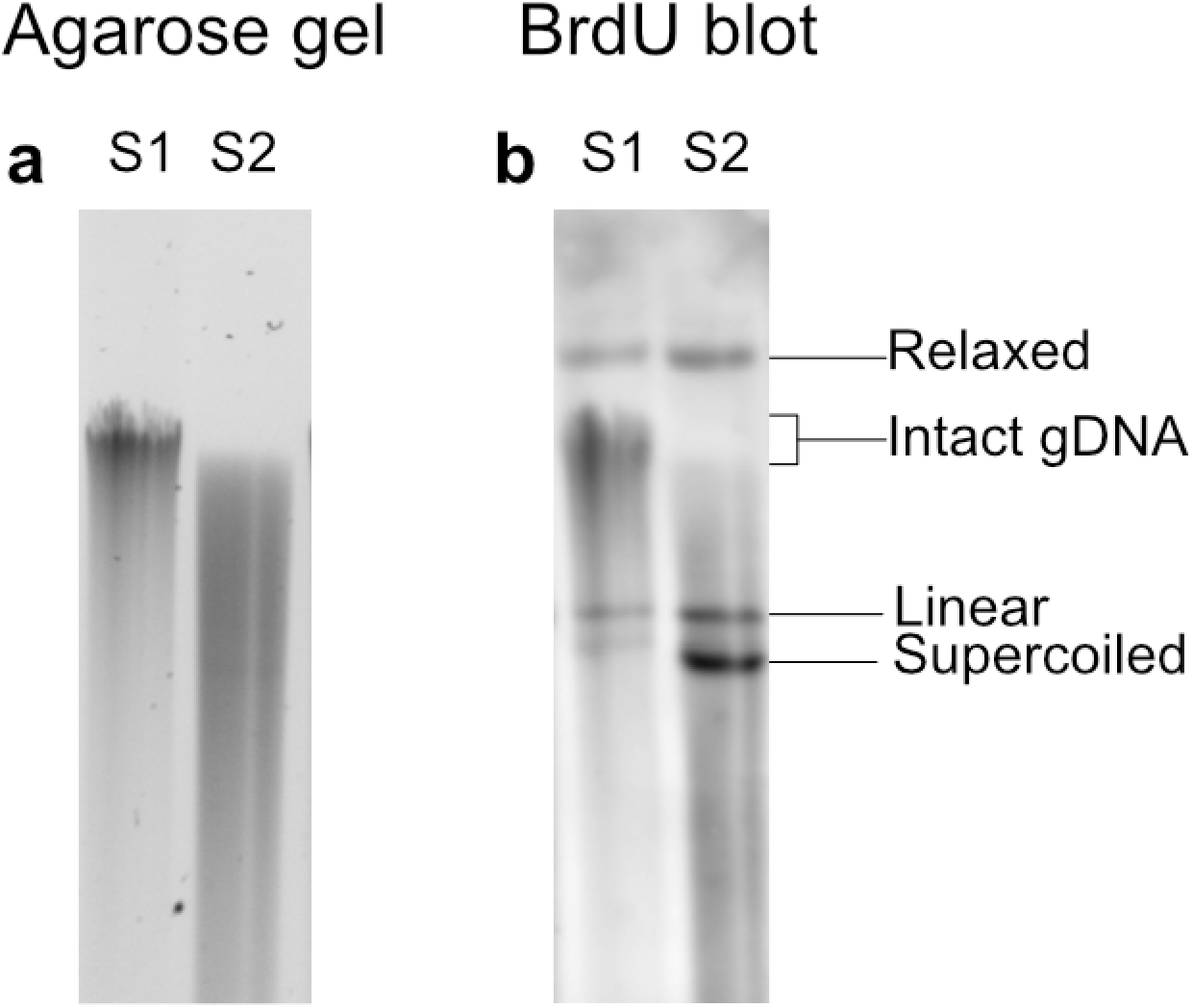
Background signal from undigested nuclear DNA. 1 µg of total DNA was isolated from HCT116 cells, treated as indicated, and separated on an agarose gel (**a**) with BrdU visualized after transfer and immunoblotting (**b**). DNA isolation was completed per manufacturer’s recommendations (S1) or with the suggested modifications (S2).

### 3.8 mtDNA supercoiling assay (Figure 3)

1. Seed cells in a 10 cm cell culture plate (*see* **Note 3**).
2. Proceed with steps 2 and 3 of section 3.1 to allow cells to become near-confluent, block nuclear DNA synthesis, and initiate BrdU incorporation. In step 4, allow BrdU incorporation for 24h.
3. Isolate DNA as described under 3.2. For samples where mtDNA topology is to be investigated, do not digest the DNA with restriction enzymes, as linearizing mtDNA will remove its topology.
4. Optional: Before proceeding to section 3.4, control samples for relaxed and linear mtDNA bands can be generated. To generate a positive control for relaxed mtDNA (Figure 3, lane 4), incubate 1 µg of DNA with topoisomerase I as recommended by the manufacturer. To generate a control for linear mtDNA fragments (Figure 3-lane 3), a restriction enzyme digest that cuts the mtDNA once (e.g., *Bam*HI for human mtDNA) can be used as described in 3.3.
5. During the agarose gel run in section 3.4, load 1 µg of DNA and use TBE buffer rather than TAE buffer and run the gel for 16-18 h at 30 V at 4 °C (*see* **Note 14**).
6. Proceed through sections 3.3–3.5 and image the final BrdU immunoblot (Figure 3b)

## 4. Notes

1. Although the supplier recommends use of fresh aphidicolin solutions, we have stored aliquots for up to 6 months at −80 °C without noticeable loss in function.
2. 5-ethynyl-2′-deoxyuridine (EdU) can be used as an alternative to BrdU. However, EdU requires different methods to visualize (*e.g.*, EdU antibody or Click-chemistry [28]).
3. Cell seeding numbers depend on cell type and desired outcomes and may need to be adjusted. In general, for assays that require imaging of a single linear mtDNA, a near confluent well of a 6-well (35 mm well diameter) is sufficient to yield the recommended 0.5 µg of DNA. To examine mtDNA topology, more mtDNA is required (at least 1 µg), and it is recommended to use a confluent 10 cm dish. This protocol has been successfully applied to HCT116 cells, human fibroblasts cells, and U2OS cells which are seeded at 400,000 cells, 450,000 cells, and 400,000 cells per well of a 6-well plate (35 mm well diameter) or 2,000,000 cells, 2,500,000 cells, and 2,000,000 cells of a 10 cm dish respectively.
4. Depending on the cell type, allowing the cells to grow to near confluence may help reduce nuclear DNA replication, which can overwhelm the mtDNA signal when cells are pulsed with BrdU.
5. Aphidicolin is added to reduce nuclear background (Figure 6). It must be added for at least 4 h prior to BrdU but can be added for longer periods (up to 24 h depending on assay timing). In cases where aphidicolin treatment is undesirable, alternative methods can be used to reduce nuclear background or enrich for mtDNA. For example, prior to DNA isolation, mitochondria can be enriched via subcellular fractionation [29,30] or isolation using a commercial kit [31]. Alternatively, mtDNA can be enriched during DNA isolation by using a miniprep kit [32, 33].
6. A minimum elution volume helps to ensure DNA concentration is sufficiently high (at least for subsequent steps, as the yield of DNA may be low for certain cell types).
7. The restriction enzyme choice will depend on the species of the cells being examined (Figure 7). *Bam*HI can be used for human DNA to resolve a single linear band for synthesis or turnover assays. However other restriction enzymes can be used to generate mtDNA fragments of various sizes. Note that the observed restriction digest pattern may vary based on cell type and mtDNA haplotype.
8. 0.5 µg of DNA per reaction is recommended when aiming to visualize a single linear band of mtDNA when using a 9 cm × 11 cm agarose gel with 10 1.5 mm wells (approximate volume 53 µL). However, for visualization of mtDNA topology, or other times when multiple bands are expected, increasing DNA to 1 µg increases the signal sufficiently to visualize multiple bands.
9. Many different loading dyes/DNA dyes can be used for the agarose gel electrophoresis. However, there are important considerations for staining and imaging the agarose gel that are relevant to the supercoiling assay (Step 3.8). First, it is recommended to stain the gel with ethidium bromide or similar DNA stain after the gel has been run, as DNA dye added the to the loading buffer or the gel can intercalate with mtDNA and affect its topology and migration through the gel. It is also important that UV imaging of the gel is done only once at the end of the assay, as UV imaging of the agarose gel mid-way through the runtime to check separation alters the conformation of mtDNA during the supercoiling assay (Figure 8).
10. Any appropriately sized object can work here, as long as it is large enough to support the agarose gel and elevates at least 3 cm above the 10X SSC buffer.
11. Depending on the imaging system, there may be an option to optimize contrast for a region of interest. This may be helpful to enhance the signal from the mtDNA bands compared to the nuclear background.
12. Although it is possible to freeze cell pellets and store them prior to DNA isolation, it is recommended to isolate DNA from fresh cells to have the best yield and preserve mtDNA topology, especially when looking at mtDNA supercoiling.
13. The high concentration of uridine is required to outcompete incorporation of residual BrdU into mtDNA that is newly replicated during the chase period.
14. The supercoiling assay requires greater separation of the similarly sized relaxed and supercoiled mtDNA topoisomers. As a result, it is recommended to running the gel overnight at a lower voltage with TBE buffer rather than TAE due to TBE’s greater buffering capacity.

**Figure 6:**
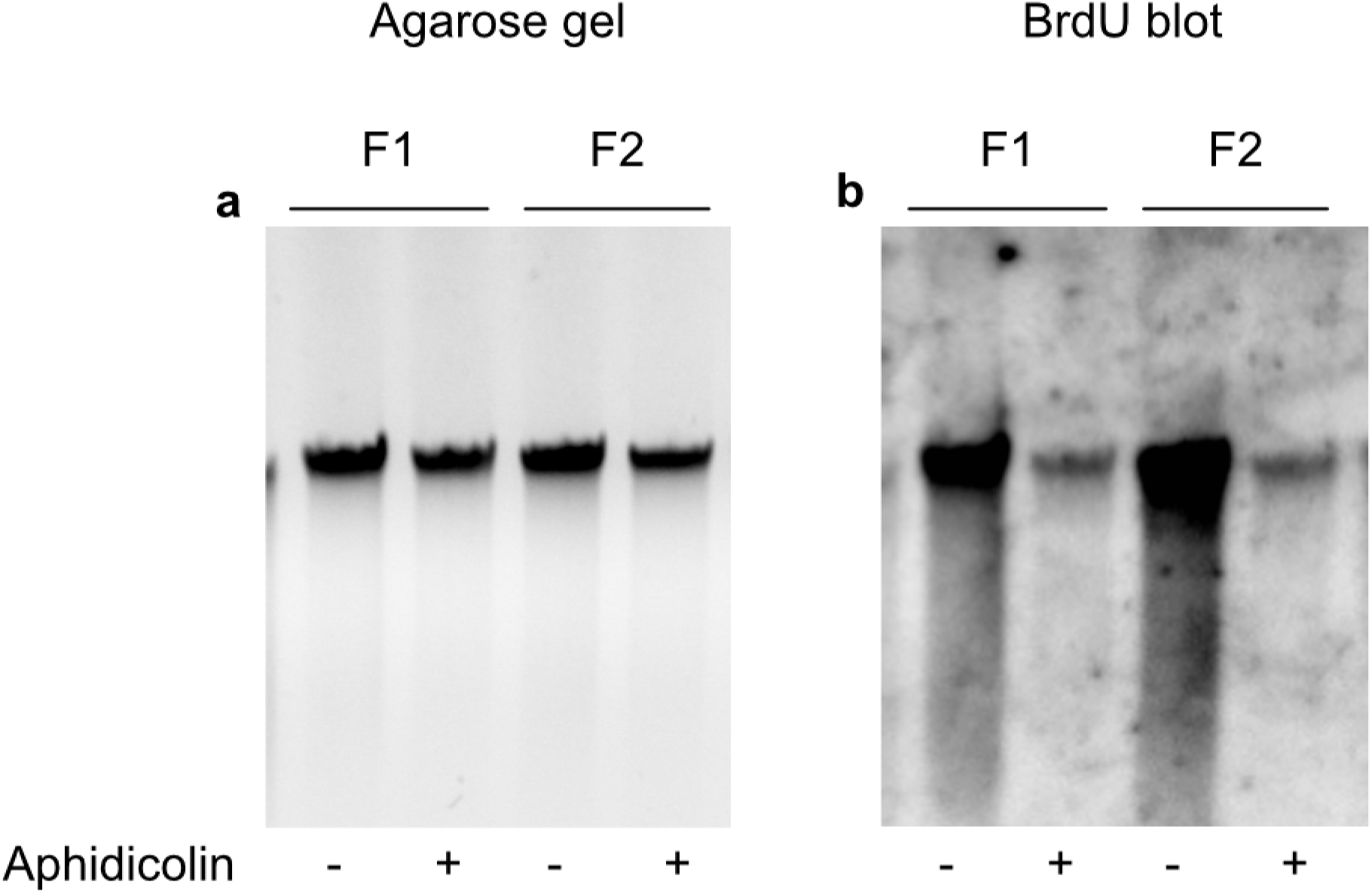
Aphidicolin reduces nuclear DNA background. 0.5 µg of total undigested DNA isolated from two primary human fibroblast lines (F1 and F2) was treated with aphidicolin as indicated (10 µM for 4 h prior to BrdU incorporation and 10 µM during BrdU incorporation for 4 h) and separated on an agarose gel (**a**). BrdU-labelled DNA was visualized after transfer and immunoblotting (**b**).

**Figure 7:**
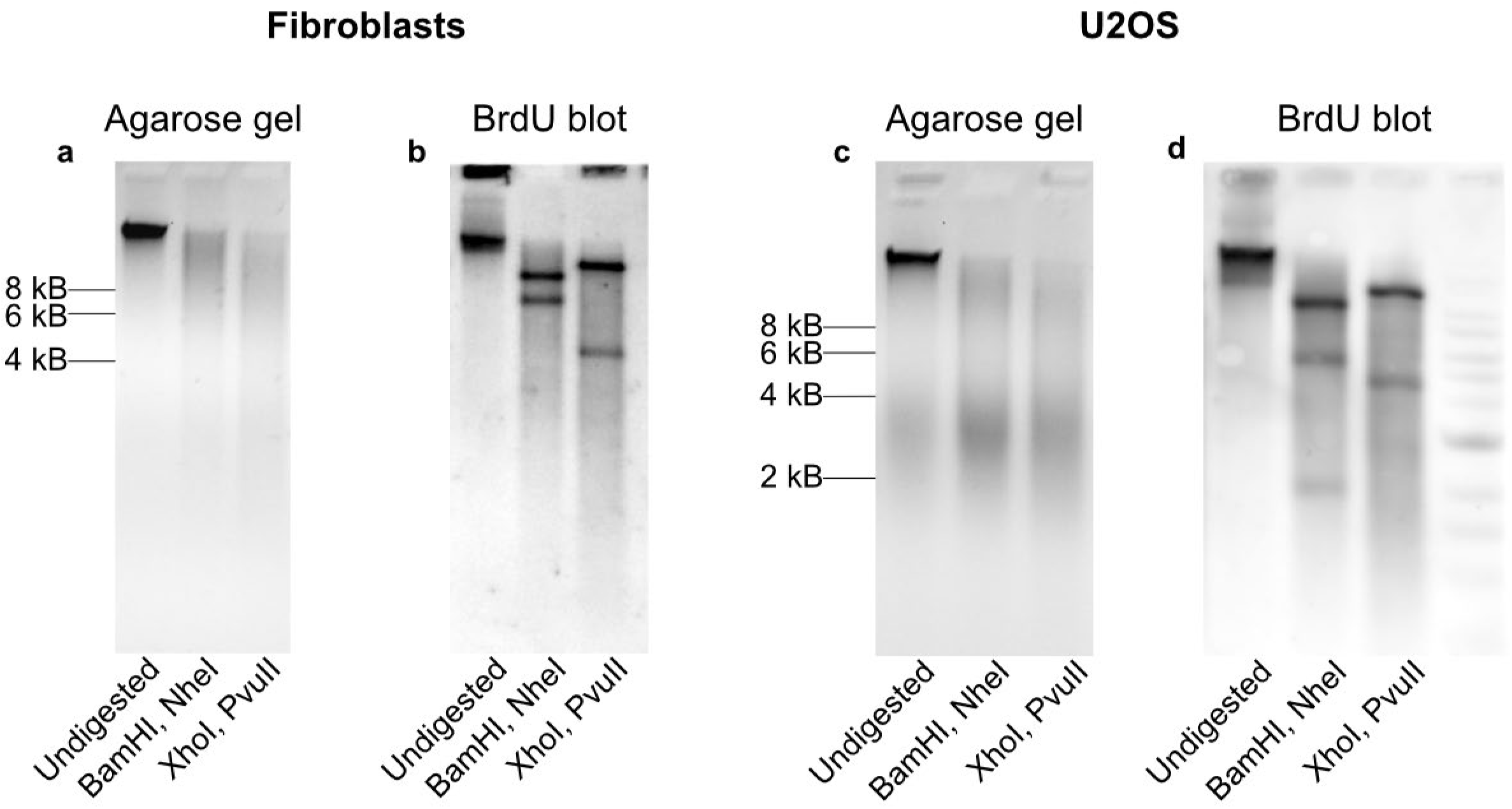
Alternative restriction enzyme digests for human mtDNA. 0.5 µg of total DNA isolated from U2OS cells or human fibroblasts was treated as indicated, separated on an agarose gel (**a, b**) and BrdU visualized after transfer and immunoblotting (**c, d**) illustrating undigested mtDNA, double digest with *Bam*HI and *Nhe*I, or double digest with *Xho*I and PvuII. mtDNA from U2OS cells shows an extra band following *Bam*HI and *Nhe*I double digest around 2 kB, likely due to differences in mtDNA haplotype

**Figure 8:**
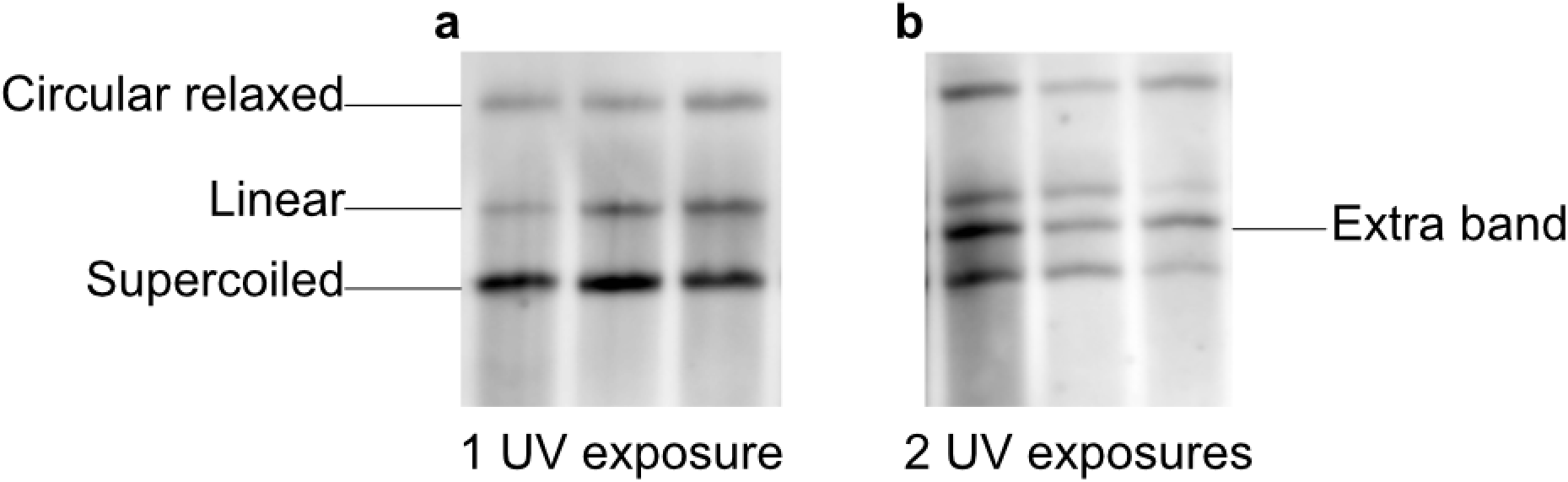
Multiple UV exposures compromise mtDNA supercoiling assay. mtDNA supercoiling assay was performed on three different HCT116 samples. 0.5 µg of total DNA isolated from HCT116 cells was treated as indicated, separated on an agarose gel and BrdU visualized after transfer and immunoblotting. BrdU immunoblots are shown from agarose gels that were imaged once at the end of running (**a**) or imaged once mid-way through running, then resumed (**b**). Extra UV exposure leads to the creation of an extra band, and alteration of mtDNA topology.

## Notes

### Competing Interest Statement

The authors have declared no competing interest.

### Summary of Updates

Correcting the recipe for TBST, which should be 0.1% Tween instead of 1% Tween.

### Funding

- Natural Sciences and Engineering Research Council

